# Steamboat: Attention-based multiscale delineation of cellular interactions in tissues

**DOI:** 10.1101/2025.04.06.647437

**Authors:** Shaoheng Liang, Junjie Tang, Guanghan Wang, Jian Ma

**Author notes:** Correspondence (J.M.) and (S.L.).

## Abstract

Spatial-omics technologies profile cells in their native spatial context within tissues, enabling more complete understanding of cellular properties. However, a key computational challenge remains: identifying cellular interactions that underlie cell types and states – interactions that are essential for spatial organization and provide a biologically grounded framework for understanding cell identities and spatial patterns. These interactions span different distances and thus require multiscale modeling, which remains a major gap in existing methods. Here, we introduce Steamboat, an interpretable machine learning framework that leverage a self-supervised, multi-head attention model to uniquely decompose gene expression of a cell into multiple key factors: intrinsic cell programs, neighboring cell communication, and long-range interactions. By applying Steamboat to diverse tissues in health and disease across various spatial-omics technologies, we demonstrate its ability to uncover critical multiscale cellular interactions, capturing classical contact signaling and revealing previously unrecognized patterns of cellular communication. Steamboat provides a powerful approach for spatial-omics analysis, offering new insights into the multiscale spatial organization of cells and their communication across a wide range of biological contexts.

## Introduction

The recent development of spatial-omics technologies has greatly enhanced our ability to study cellular states within their native spatial environment, providing key insights into diseases and developmental processes [1–3]. For instance, spatial transcriptomics has been instrumental in characterizing immune responses from tertiary lymphoid structures in cancer [4] and mapping dysfunctional cellular communications in brain demyelination [5]. Large-scale initiatives such as the Human Cell Atlas [6], the BRAIN Initiative [7, 8], and the Human Tumor Atlas Network [9] have further propelled research by generating comprehensive spatial cellular atlases [10–12], facilitating a deeper understanding of tissue organization and function. At the core of these analyses lies the critical role of cell-cell interactions, which are fundamental to maintaining homeostasis and orchestrating multicellular development [13, 14]. The ability to examine these nuanced intercellular interactions in spatially resolved contexts is essential for decoding the complex cellular communication patterns and dynamics that govern health and disease [1, 13].

Characterizing cell-cell interactions has garnered significant attention [13]. However, many existing approaches rely on ligand-receptor databases [15–17], which are incomplete [13] and often miss tissue-specific interactions. Other methods assume a fixed number of neighboring cells constitute an “interacting neighborhood”, or use fixed radius or Delaunay triangulation-based approaches [18–21], overlooking the fact that not all nearby cells interact meaningfully. A clear example is found in the intestinal epithelium, where tight junctions effectively prevent molecular exchange, cutting off direct communication [22]. Furthermore, cellular interaction varies in strength, as seen in the dynamic signaling between T cells and dendritic cells during antigen presentation [23]. While models like NCEM [24] can infer interaction strengths, they depend on accurate cell-type annotations. SIMVI [25] can disentangle intrinsic and spatially induced gene expression without annotation, but its graph attention network is restricted to short-range interactions. DECIPHER [26] generates spatial context encoding independent of the center cell, failing to capture cell type-specific interactions. Crucially, none of these methods models multiscale interactions, particularly missing long-range dependencies. A method capable of embedding cells within their context while quantifying multiscale cell-cell interactions *de novo* from spatial-omics data remains unavailable.

To address these limitations, we develop Steamboat, an interpretable, self-supervised, multi-head attention model for revealing multiscale cellular interactions. Inspired by signaling mechanisms across multiple scales – such as direct contact and secreted signaling molecules [27] – Steamboat decomposes spatial transcriptomic profiles into three spatial scales: (1) intrinsic (ego) effects, (2) micro-environmental (local) effects, and (3) sample-wide macro-environmental (global) effects. This decomposition yields three key outputs: cell embeddings, intercellular interaction graphs, and environment-aware reconstructed cells. The interaction graph in Steamboat enables various types of analyses related to cell-cell communications, including ligand-receptor interactions. Its reconstruction mechanism uniquely enables *in silico* spatial perturbation, allowing the prediction of a cell’s response to a changing environment. We demonstrate that Steamboat’s cell-cell interaction-aware approach outperforms interaction-agnostic spatial smoothing methods [28–30] for cell type clustering and spatial domain segmentation, which otherwise blur gene expression [31]. We further illustrate Steamboat’s potential to uncover patient-level prognostic markers and its unique capability for spatial perturbation analysis, establishing it as a powerful tool for spatial-omics analysis, particularly in decoding multiscale cellular interactions.

## Results

### Overview of attention-based multiscale modeling in Steamboat

At the core of Steamboat is its ability to decompose a cell’s molecular profile (e.g., gene expression) into three contributing spatial scales: ego, local, and global – corresponding to intrinsic gene programs, the microenvironment, and the macroenvironment, respectively (**Fig. 1a**). This is achieved using a novel design of the multi-head attention mechanism [32] in Steamboat, tailored for spatial-omics data (**Fig. 1b**).

**Figure 1.**
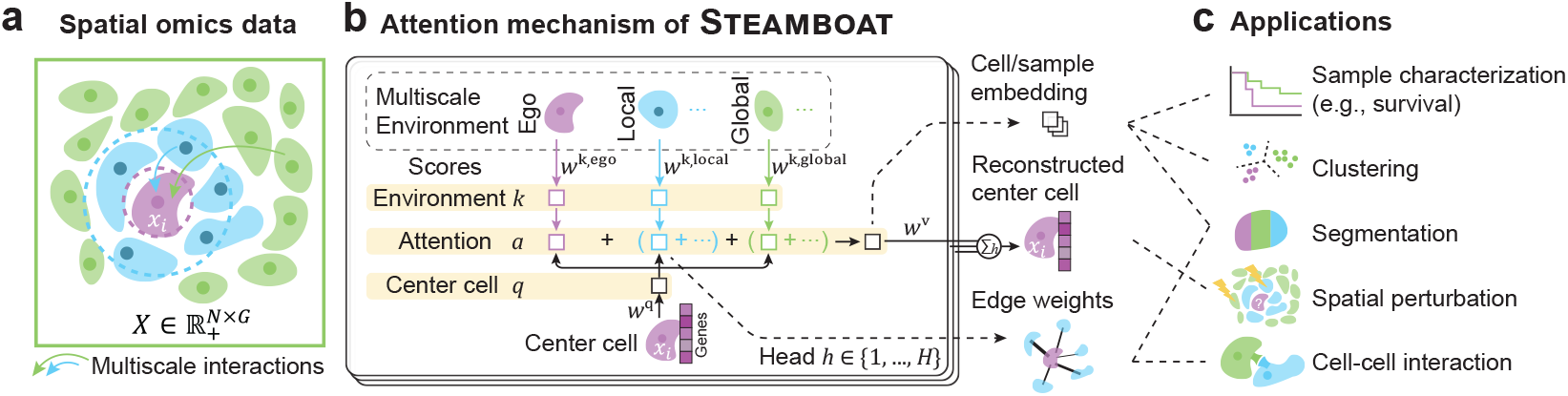
Overview of the Steamboat framework. **a**. The input to Steamboat consists of gene expression (or other molecular profiles) of cells and their spatial coordinates. The environment around each cell is decomposed into three scales: ego (purple), local (blue), and global (green). Arrows represents interactions at different scales. **b**. Steamboat optimizes five metagenes: one for the center cell, three for the cells it interacts with at the ego, local, and global scales, and one reconstruction metagene. The model minimizes the reconstruction error of the center cell. Attention between two cells depends on the center cell’s score *q*, the environment score *k* of the other cell, and the distance between the cells. Attention scores between cells form a weighted cellular interaction graph, and the total attention score of each cell is used as a cell embedding, which is then used to reconstruct the center cell. **c**. These embeddings and interaction graphs enable a variety of downstream tasks, including spatial domain segmentation, cell type clustering, *de novo* cell-cell interaction discovery, spatial perturbation, and patient survival prediction. Dotted lines between panels **b** and **c** indicate the specific components of Steamboat used in each application.

Steamboat employs the attention mechanism to recognize cell-cell interactions, which vary across cell types and states. To disentangle different interactions, multiple “attention heads” are used. The term “head” is borrowed from natural language processing, where it originally refers to a submodule trained to detect specific interactions between words [32] (see **Methods** for details). Each head is parameterized by metagenes, which are weighted combinations of signature genes associated with cell types or biological functions [33, 34]. Two cells are considered to interact under a given attention head if one is selected as a center cell and the other as part of the interacting environment. Mathematically, this selection is performed via an inner product between a cell’s gene expression profile and the metagene, effectively comparing a cell to a learned cell signature and producing center cell scores and environment scores. (There is no hard threshold to actually select a cell – only real-valued scores.) We quantify interaction strength using the product of the center cell score and environment score, referred to as the attention score.

Steamboat dissects attention into three spatial scales: global, local, and ego, each with its own metagene. Global attention captures a cell’s interaction with the broader tissue context (e.g., secreted signaling molecules), while local attention captures spatially proximal interactions (e.g., direct cell contact). A center cell may lack strong dependencies at these levels; thus, an ego scale is introduced to allow a cell to attend to itself. Attention scores are summarized across heads to construct a cell embedding, while local attention scores are used to build a weighted cellular interaction graph (**Fig. 1b**). The weights of genes in these metagenes are optimized in an unsupervised manner by reconstructing the center cell from the cell embedding using reconstruction metagenes. In total, each head uses five metagenes: one for the center cell, three for the three spatial environment scales, and one for reconstruction.

Disentangling attention across spatial scales is crucial for capturing biologically meaningful interactions. Traditional attention mechanisms are often not directly interpretable [35] and may overfit to spurious long-range dependencies [36]. Design choices in Steamboat, such as enforcing nonnegative weights, further enhance interpretability.

In summary, Steamboat produces three key outputs: a cell embedding, a weighted cellular interaction graph for each attention head, and a reconstructed center cell integrating both self (intrinsic) and environmental information. These outputs support a range of downstream analyses for understanding spatial interactions within tissues (**Fig. 1c**). Further details of the Steamboat model can be found in the **Methods** section.

### Simulation reveals interpretable outputs of Steamboat

To illustrate the outputs of Steamboat, we designed a simple simulation with two layers of cells (**Fig. 2a**). The first layer consists entirely of cell type A, while the second layer contains a random mixture of cell types B, C, and D. Each cell type is characterized by two marker genes (e.g., *A1-2* for cell type A). To simulate intercellular interaction, cell types B and C express the receptor gene *R* when adjacent to cell type A.

**Figure 2.**
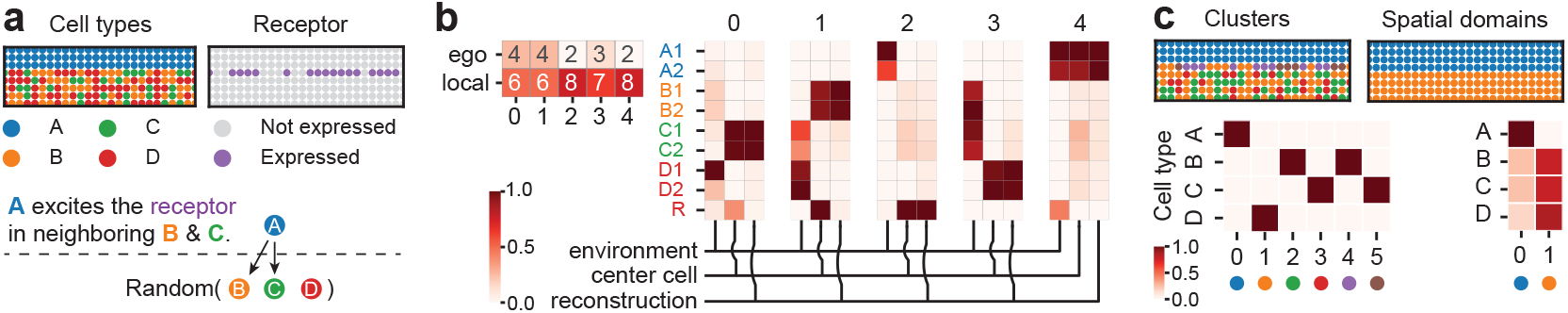
Simulation illustration of Steamboat. **a**. Spatial distribution of cell types and receptor expression in the illustrative simulation. The upper region exclusively contains cell type A, while the lower region includes a random mixture of cell types B, C, and D. The receptor is expressed in B and C cells that are adjacent to A cells. The spatial graphs show a subset of the dataset, with this pattern repeating horizontally. **b**. Evironment, center cell, and reconstruction metagenes across all attention heads. The top left corner indicates the relative importance of ego and local environments. Gene names are color-coded to correspond to the receptor and cell types in panel **(a). c**. Clustering and spatial domain segmentation results from Steamboat, with accompanying confusion matrices comparing these results to the ground truth cell types.

Using five attention heads, Steamboat decomposes these interactions into five sets of metagenes, each consisting of environment, center cell, and reconstruction metagenes (**Fig. 2b**). The interpretation is intuitive: in an environment defined by the environment metagene (*k*), if the center cell expresses center cell metagene (*q*), it will also express metagene (*v*).

In this example, head #0, #1, #3, and #4 correspond to the four cell types C, B, D, and A, respectively. Notably, the environment metagene in head #4 indicates that cell type A is most likely to neighbor another A cell, while heads #0, #1, and #3 suggeset that cell types B, C, and D are colocalized. Importantly, head #2 highlights signaling from cells expressing *A1-2* to those expressing *B1-2, C1-2*, and *R*, capturing the simulated signaling pathway. We then calculated the relative importance in reconstruction (**Fig. 2b**), and the local attention scale is the highest in head #2, further demonstrating Steamboat’s ability to differentiate intrinsic programs and environmental influences. Head #4 is also high in local attention, likely due to the concentrated spatial distribution of cell type A. Clustering and spatial domain segmentation based on Steamboat embeddings and cell-cell network construction closely align with the simulated cell types and spatial regions (**Fig. 2c**).

Together, this simulation demonstrates the interpretable mechanisms uncovered by Steamboat’s attention-based framework.

### Steamboat characterizes cell-cell interactions in ovarian cancer

To examine Steamboat’s ability to embed cells and characterize cell-cell interaction (CCI), we applied it to cancer data, where CCI has been widely studied. Yeh et al. [37] introduced a comprehensive dataset of tubo-ovarian high-grade serous carcinoma (HGSC), aiming to elucidate mechanisms of immune evasion and therapeutic vulnerabilities. Here, we focus on a more homogeneous subset of this dataset – 27 CosMx untreated adnexa samples – each containing 2,000 to 10,000 cells with 979 measured genes (e.g., **Fig. 3a**) is used. We applied 25 attention heads to decompose gene expression (see Number of attention heads in **Methods**), which automatically attend to ego, local, and global environments.

**Figure 3.**
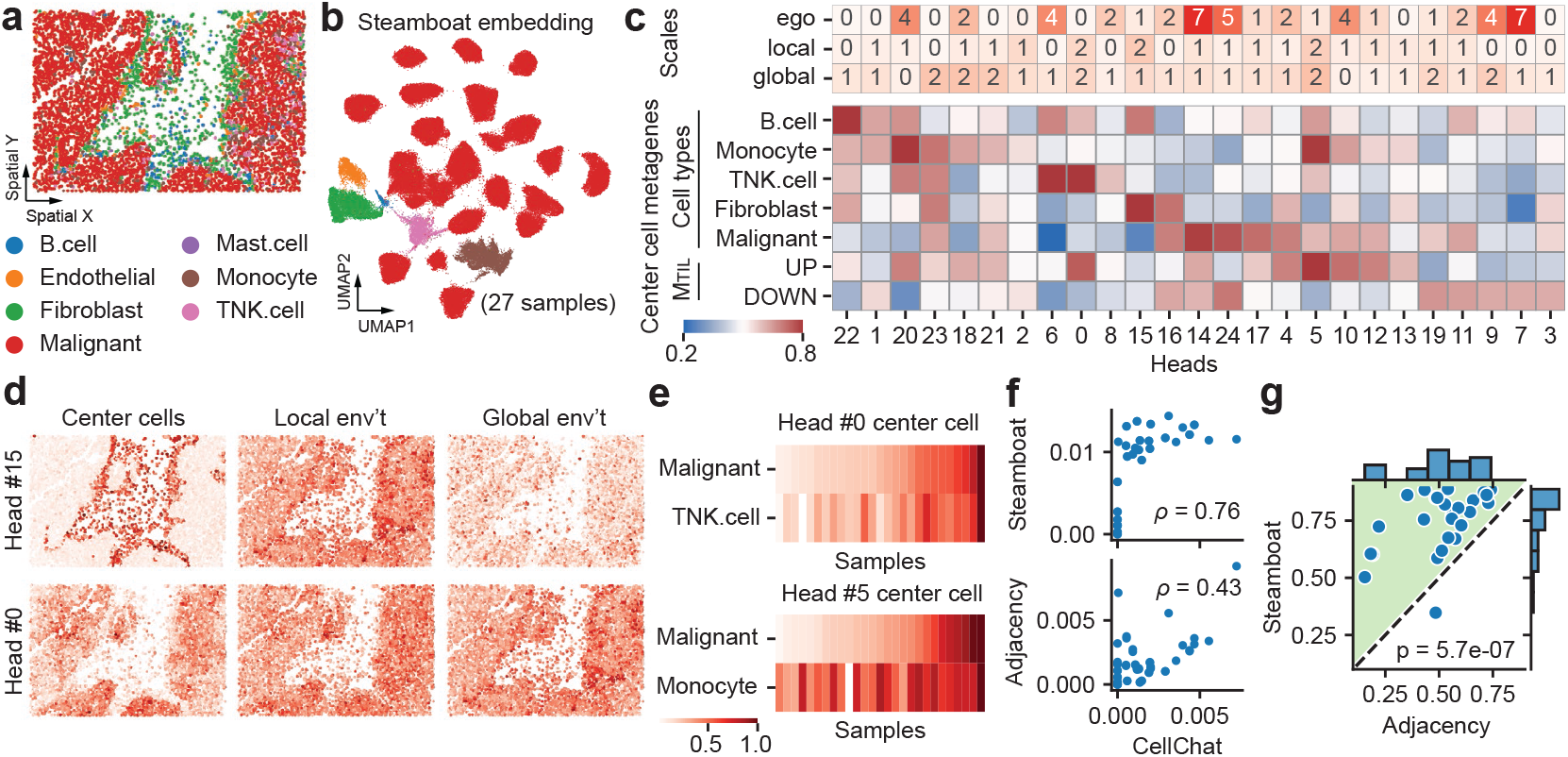
Steamboat characterization of ovarian cancer samples. **a**. Spatial distribution of cell types of a representative slide (SMI_T10_F001). **b**. UMAP visualization of Steamboat embedding, colored by cell types (see **Fig**. S1a for the same UMAP colored by sample). **c**. Top: Weight of each head and spatial scale, normalized to a total of 100. Bottom: Gene set enrichment of Steamboat center cell metagenes (*>* 0.5: positive enrichment, *<* 0.5: negative enrichment; see **Methods**). See **Fig**. S1b for enrichment of other metagenes). **(d)** Spatial distribution of cells, colored by Steamboat metagene scores. **e**. Average center cell scores for malignant cells, T/NK cells, and monocytes across all samples. **f**. Comparison of cell-cell interaction scored from Steamboat, adjacency matrix, and CellChat on sample SMI_T10_F001. Each dot represents a pair of cell types. Spearman correlations (*ρ*) measure agreement between methods. **g**. Summary of comparisons across all 27 samples. *P*-value from Mann-Whitney U test.

We first examined the embedding and metagenes generated by Steamboat. UMAP visualization of the embedding shows that Steamboat accurately captures cell types, as well as the patient-specific cancer cells (**Fig. 3b** and **Fig**. S1a). To interpret Steamboat’s attention heads, we calculated the enrichment of cell type-specific gene sets (from the original pulication [37]) in Steamboat metagenes (**Fig. 3c** and Fig. S1b). As expected, each cell type is represented by a few heads (**Fig. 3c**) – for example, head #6 and #0 for T/NK cells, #15 and #16 for fibroblasts, and #14, #24, #17, and #4 for malignant cells. The clear clusters in the UMAP and the cell type-specific attention heads demonstrate that Steamboat faithfully represents the gene expression profiles of the center cells.

We also calculated the contribution of the three scales in every head (**Fig. 3c**). For exmaple, head #15 selects fibroblasts as center cells (**Fig. 3c,d**) and reveals a local (short-range) interaction with other fibroblasts (**Fig**. S1b). It also detects a global interaction between fibroblasts and B cells (**Fig. 3c** and **Fig**. S1b), which are known to interact via long-range cytokine signaling [38]. Among the top 20 genes of the global environment metagene, eight are related to signaling receptor binding, and three to cytokine binding (**Fig**. S1d) [39]. As another example, head #0 shows interactions between T/NK cells and cells expressing genes promoting tumor immune infiltration, discussed in the next paragraph. Interaction patterns are observed in other heads as well (e.g., **Fig**. S1c).

The original publication discovered M_TIL_ (malignant TIL), a transcriptional program in malignant cells that marks the presence of TILs. M_TIL_ consists of 100 upregulated and 100 downregulated genes that best predict T/NK cell infiltration in a supervised learning setting. It also provides gene sets for cell type signatures. Heads #0 and #5 are both enriched for M_TIL_-up (**Fig. 3c**). Interestingly, these heads primarily select T/NK cells and monocytes, not malignant cells, although malignant cells in some samples received high scores (**Fig. 3e**). In contrast, heads primarily selecting malignant cells (#14, #24) are all enriched in M_TIL_-down (**Fig. 3c**). This suggests that while cancer cells generally inhibit TIL, some samples contain malignant cells that express genes similar to T/NK cells and monocytes that promote TIL. Supporting this interpretation, there is significant overlap (*p* = 3 × 10^*−*6^, hypergeometic test) between two gene sets – one in monocytes and one in malignant cells – both associated with promoting TIL, as reported in the original study. Other work has simimilarly found chemokine secretion by both malignant cells and monocytes in various cancers, including ovarian cancer [40, 41]. To further investigate how these cells interact with others, we examined the local and global environment metagenes (**Fig**. S1b), which show interactions with B cells, monocytes, and T/NK cells – consistent with the M_TIL_-up signatures.

To more comprehensively assess Steamboat’s ability to recognize CCI and test the difference between cell proximity and functional interaction, we compared Steamboat with other CCI inference methods. To the best of our knowledge, no definitive ground truth for CCI exists. However, ligand-receptor databases (LRDBs) are commonly used to infer CCI in scRNA-seq and spatial omics data [42–44], with CellChat [43] providing the most support for spatial omics data. We used CellChat to calculate interaction strengths between cell types, and summarized Steamboat’s cell-cell attention scores by cell types to produce a CCI matrix for each sample. For comparison, we also computed proximity-based interactions by counting the pairs of adjacent cells between cell types, normalized by cell numbers. Strikingly, Steamboat’s CCI matrices strongly agree with those from CellChat in a representative sample (SMI_T10_F001) (Spearman *ρ* = 0.76, **Fig. 3f**). In contrast, proximity-based inference (**Methods**) shows much lower agreement (*ρ* = 0.43), highlighting the distinction between spatial proximity and true interactions. This pattern holds across all samples (**Fig. 3g**). While LRDBs are not comprehensive, the high agreement with CellChat demonstrates that Steamboat – a purely data-driven method that does not rely on LRDBs – can capture true CCI signal *de novo*.

Together, these findings show that Steamboat effectively recognizes cell types, states, and interactions.

### Steamboat identifies cell types and spatial domains in mouse brain

Clustering and spatial domain recognition are key tasks in spatial omics studies. Here, we demonstrate how Steamboat unifies these two tasks within a single, cohesive framework, using the whole mouse brain atlas assayed by MERFISH [10]. We also illustrate its capability for spatial perturbation and ligand-receptor interaction recognition. The dataset includes 129 adult mouse brain slides with 2.6 million cells, profiled across a 1,122-gene panel. The data are annotated with 35 major cell types (classes; **Fig. 4a**) and various brain anatomical regions (**Fig. 4b**), as provided by the Allen Institute, offering valuable insights into brain organization, cellular diversity, and the roles of different cell types in brain function and development [10].

**Figure 4.**
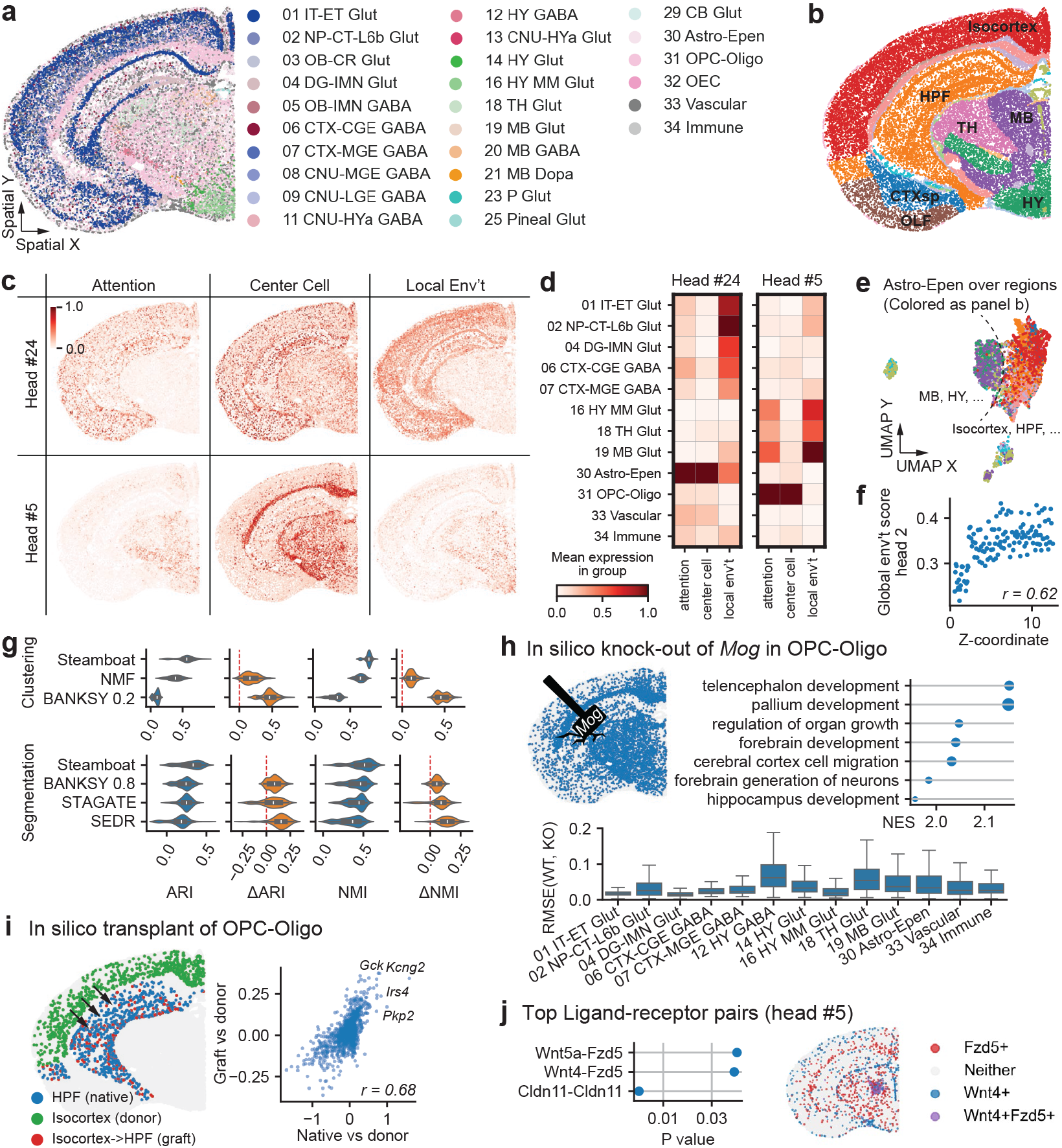
Mouse brain MERFISH data analysis. **a-b**. Representative slide showing cell types **(a)** and spatial domains **(b)**; note that not all cell types and spatial domains are present in this slide. Refer to the **Supplementary Notes** for a list of abbreviations. **c**. Scores of cells assigned by two attention heads #24 and #5. **d**. Cell scores for each metagene, summarized by cell types. Only cell types comprising >1% of total cells are shown. **e** UMAP of Astro-Epen cells, colored by brain region. **f**. Correlation between the global environment score of head #2 and the z-coordinate of the slides. **g**. ARI and NMI scores of Steamboat and other methods across all slides, for spatial domain segmentation and cell type clustering. Columns 1 and 3 show overall ARI and NMI distribution; columns 2 and 4 show per-sample improvements of Steamboat over other methods. Steamboat is significantly better in all comparisons (paired Wilcoxon signed-rank test). **h**. Knock-out scenario of spatial perturbation. Top-left: schematic illustrating knock out of *Mog* in OPC-Oligo cells. Top-right: top pathways changed in HY GABA, identified by GSEA. Bottom: magnitude of expression changes in non-OPC-Oligo cell types. **i**. Transplant scenario in spatial perturbation. Left: schematic showing transplantation of OPC-Oligo cells from the isocortex to the HPF. Right: correlation of log-fold changes between graft vs. donor and native vs. donor comparisons. **j**. Enriched ligand-receptor pairs in head #5, and spatial distribution of *Wnt4* and *Fzd5*.

We trained Steamboat on all samples using 50 attention heads (**Methods**) Notably, head #24 assigns markedly different center cell scores and attention scores to cells (**Fig. 4c**), indicating strong environmental influence. While the center cell metagene selects all astrocytes and ependymal cells (Astro-Epen), the local environment scores are high only in neurons located in the isocortex and hippocampal formation (HPF) (**Fig. 4d**), thereby selecting Astro-Epen in these regions only. In fact, there is a clear difference in the gene expression of Astro-Epen in different regions (**Fig. 4e**). Similarly, in head #5, the attention mechanism distinguishes OPC and oligodendrocytes (OPC-oligo) in the thalamus, hypothalamus, and midbrain from those in other regions (**Fig. 4c,d**), again showing Steamboat’s ability to distinguish cell groups using both the gene expression of the center cell and its spatial neighborhood.

Because the samples in this dataset come from the same donor, differences between slides are small, resulting in generally low global attention (**Fig**. S2a). Nevertheless, in head #2 (the one with the highest global attention), the global environment score is clearly correlated to the z-coordinates of the samples (**Fig. 4f**). These results demonstrate that Steamboat’s multi-scale attention can represent rich information in the data.

To compare Steamboat with other methods, we used the annotated spatial domains (“division” in the original publication [10]) as reference and calculate ARI and NMI for each slide. Steamboat outperforms BANKSY [30], STAGATE [28], and SEDR [20] in spatial domain segmentation, in both pooled and per-sample comparisons (**Fig. 4g** and **Fig**. S3). An ablation study substituting Steamboat’s local attention graph with a spatial *k*-NN graph (**Fig**. S2b) results in decreased performance, while the baseline of PCA with spatial *k*-NN graph (studied by [45]) performs similarly to the ablation – ruling out any incompatibility between Steamboat embeddings and *k*-NN graph. Overall, Steamboat produces a meaningful spatial graph that supports improved spatial segmentation. In cell type clustering (“class” in the original publication [10]), Steamboat also overperforms BANKSY (**Fig. 4f**). Since Steamboat can be viewed as an extension of NMF (non-negative matrix factorization) augmented with spatial interaction information, it was included as a baseline/ablation. Results show that Steamboat’s ability to model neighboring cells outperforms conventional NMF, highlighting its ability to generate informative and biologically meaningful cell embeddings.

### Steamboat enables in silico spatial perturbation and intercelluar interaction discovery

Perturbation studies are essential for understanding cellular behavior under different conditions. To overcome practical limitations of vast combinatorial perturbation experiments, *in silico* perturbations have been explored using single-cell pretrained foundation models [46, 47]. Unlike conventional *in silico* perturbation tasks, which focus on knocking out genes in a cell, Steamboat enables *in silico* spatial perturbation to predict how a cell behaves when its spatial context is altered. We explore two such scenarios: (1) how cells react to genetic perturbation in surrounding cells, and (2) how a cell behaves when placed in a different environment.

In the first scenario, we “knock out” *Mog*, which encodes myelin oligodendrocyte glycoproteinin oligodendrocytes [5], enriched in both the cortex and thalamus region (**Fig. 4h** and **Fig**. S4). Significant changes are observed not only in oligodendrocytes but also in thalamus neurons, consistent with prevalence of thalamic damage in demyelinating diseases like multiple sclerosis [48]. In oligodendrocytes, *Sox10* – a gene crucial for myelination maintenance [49] – is the second most downregulated DEGs. *Cldn11*, which encodes a major myelin component, is also among the top five. In thalamic neurons, GSEA shows changes in gene ontology sets related to neuron development, such as telencephalon development and pallium development (**Fig. 4h**), consistent with demyelination effects.

In the second scenario, we “transplant” OPC/oligodendrocytes from the isocortex to HPF. OPC is one of the most migratory cell type in the CNS [50]. UMAP shows that the gene expression of transplanted cells (“graft”) shifts toward that of OPC-oligo cells native to HPF (**Fig. 4i**). Specifically, differential expression analyses between graft vs. donor and native vs. donor show strong agreement in log-fold changes. This suggests that the transplanted cells are influenced by their new environment, adapting to resemble local counterparts – supporting the utility of Steamboat for modeling spatial perturbation.

We further used Steamboat to investigate gene programs underlying intercellular interactions with OPC-oligo cells. Among the ligand-receptor pairs curated by CellChatDB [51], 84 are present in this MERFISH panel. A permutation test identified four significant ligand-receptor pairs in head #5, with observed co-localization of cells expressing both ligand and receptor (**Fig. 4j** and **Fig**. S2c). For example, *Cldn11* appears to interact with itself, likely reflecting CCI among oligodendrocytes. *Wnt4* and *Wnt5a*, two Wnt ligands expressed in oligodendrocytes, interact with *Fzd5*, a Wnt receptor involved in neuronal survival, polarity, and neurite growth [52].

Overall, we demonstrate that Steamboat effectively reveals biological relationships between cells at multiple scales, supporting cell type clustering, spatial domain segmentation, and intercellular interaction discovery. Crucially, Steamboat uniquely enables spatial perturbation, allowing prediction of cellular responses to environmental changes. This capability can deepen our understanding of the molecular mechanisms underlying tissue organization in health and disease.

### Steamboat unveils sample-level features linked to cancer prognosis

Having shown the utility of global-scale attention in previous datasets, we next applied Steamboat to more diverse samples to further explore its potential. Specifically, we analyzed a CODEX dataset from a colorectal cancer (CRC) invasive front study involving 35 patients, divided into two risk groups: 17 CLR (Crohn’s-like reaction) and 18 DII (diffuse inflammatory infiltration) (**Fig. 5a**) [53]. CLR is characterized by the presence of tertiary lymphoid structures (TLS), also known as “follicles”, which promote lymphocyte infiltration and are associated with better prognosis (low risk) [54]. Patients without TLS were classified as DII (high-risk). Four samples were collected from each patient. For CLR patients, regions were chosen so that half of the samples contained TLS. The formalin-fixed paraffin-embedded (FFPE) samples were assayed using a reengineered CODEX method to measure 56 markers, and cell types and neighborhoods were annotated in the original study (**Fig. 5b**).

**Figure 5.**
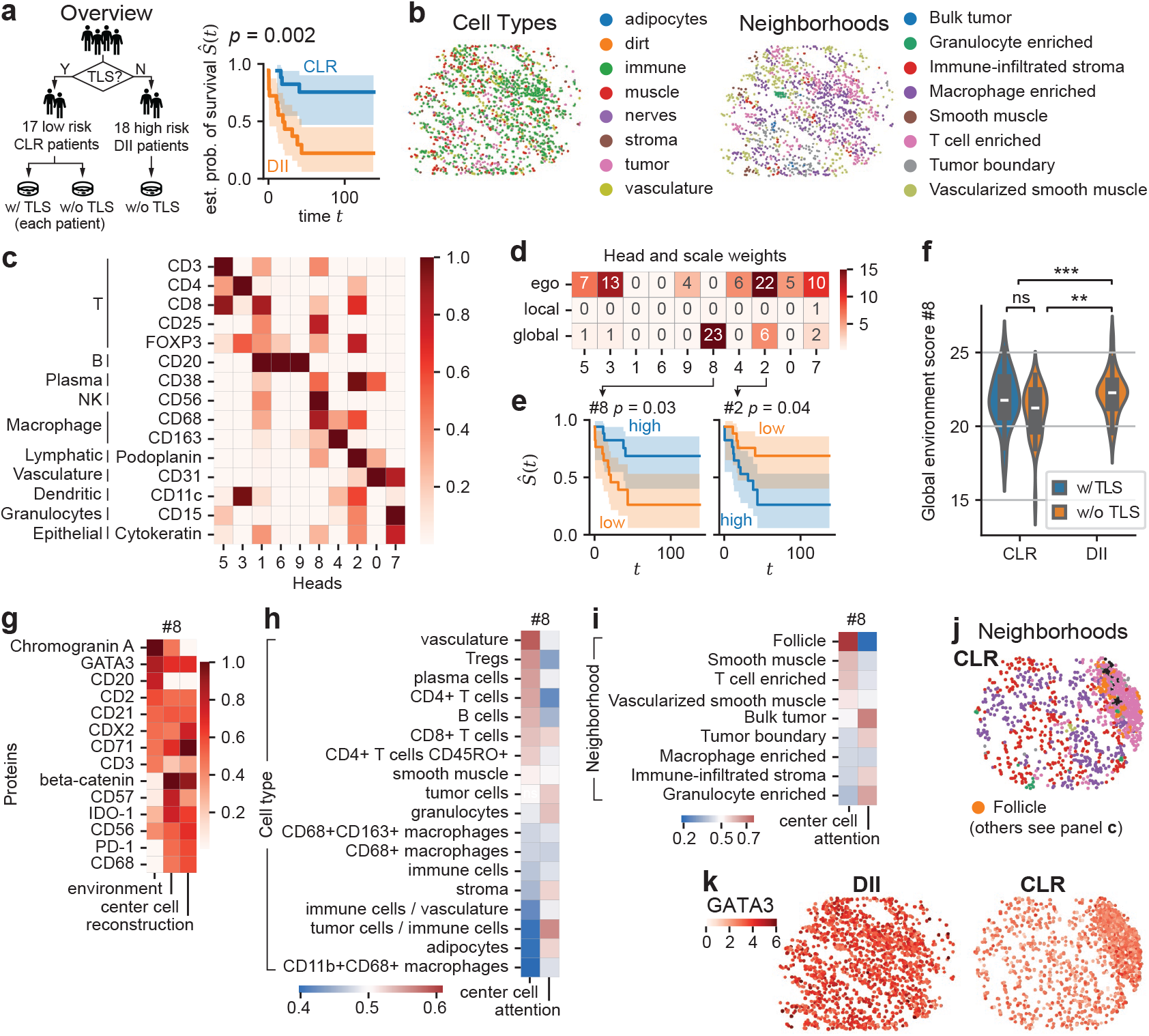
Steamboat unveils sample-level features related to colorectal cancer prognosis. **a**. Left: Experimental design from the original dataset. Patients are grouped into two risk categories based on TLS presence, with four samples per patient. DII samples are all without TLS. Samples with and without TLS are collected from each CLR patient. Right: Survival analysis (Kaplan-Meier curve and log-rank test) comparing the two clinically defined groups. **b**. Spatial distribution of cells in a representative DII sample, colored by cell types (left) and neighborhoods (right) as annotated in [53]. **c**. Center cell metagene weights for cell type markers curated from [53]. **d**. Weights of ego, local, and global attention scales across all attention heads. **e**. Survival analysis comparing patients with high (*>*median) and low (*<*median) global attention in heads #8 and #2. **f** Sample-level (global) environment scores from head #8, stratified by CLR/DII classification and TLS presence. *P*-values from Mann-Whitney U test are shown above (** *<* 0.01, *** *<* 0.001, ns: not significant) **g**. Genes with the highest loadings in the global environment, center cell, and reconstruction metagenes of head #8. **h-i**. Median center cell and attention scores of cell types (**h**) and neighborhoods (**i**) in head #8. **j-k**. Expression of GATA3 in representative CLR and DII (**k**) samples, and corresponding neighborhood annotations in the CLR sample for reference (**j**).

In this analysis, we used ten attention heads. As in previous datasets, Steamboat decomposes protein expression such that each head represents distinct cell types (**Fig. 5c**). Compared with the mouse brain dataset, the weights for global-scale attention are considerably higher, consistent with the heterogeneity among cancer samples. Among all the heads, head #8 exhibits the highest global scale attention (**Fig. 5d**), followed by head #2. To understand the environment they represent, we examined their relationship to patient and sample characteristics. Notably, attention scores of heads #8 and #2 significantly distinguish patients with different prognoses (**Fig. 5e**). For head #8, the global environment score is strongly associated with CLR/DII classification (Mann-Whitney U-test *p* = 0.0001, common-language effect size *f* = 0.66) (**Fig. 5f**). More interestingly, the score does not distinguish between TLS-present and TLS-absent CLR samples (*f* = 0.37, *p* = 0.08), yet a notable difference remains between TLS-absent CLR and DII samples (*f* = 0.71, *p* = 0.001). This suggests that the global environment metagene captures a patient-wide macroenvironment lacking TLS, rather than merely reflecting TLS absence at the sample level.

Indeed, although the center cell metagene for head #8 (**Fig. 5g-i**) selects immune cells in TLS (follicle) regions (or similar cells in DII samples), the attention is instead less on immune cells and more on tumor cells. This implies that the environment metagene represents a global environment that may act against immune cell activity. This is further supported by the genes in the metagene (**Fig. 5g**), including, for example, *GATA3*, which is globally upregulated in DII samples (**Fig. 5j,k**) and is essential for the immune-suppressive function of Tregs [55], and *CD71*, which is associated with poor anti-tumor response [56]. Head #2 (Fig. S5b) also shows a similar difference between CLR/DII groups, which have higher attention on Tregs. It highlights high expression of Ki67, a widely used tumor marker indicating proliferation [57], in the environment, and high expression of CD194 (*CCR4*) and CD5 – both markers of Treg immunosuppressive function [58] in the center cells.

This analysis demonstrates that global attention heads are sensitive to biological variation across patients and, when integrated with ego and local scales capturing specific cell types, can help elucidate communication pathways across multiple spatial scales. It also highlights Steamboat’s versatility in analyzing proteomic assays, such as CODEX, in addition to transcriptome data.

## Discussion

In this study, we introduce Steamboat, a streamlined multi-head attention model that harnesses recent advancements in neural networks with interpretability to uncover novel patterns of cellular spatial organization and dependencies. Optimized for sparse graphs, Steamboat achieves computational efficiency comparable to state-of-the-art methods, enabling practical applications across diverse spatial omics datasets.

Our applications to ovarian cancer, mouse brain, and colorectal cancer datasets demonstrate Steam-boat’s capability to uncover biological insights into intercellular dependencies across different spatial scales. Notably, global attention heads reveal proteomic patterns associated with patient prognosis, while local heads capture cell-cell interactions, such as neuron-glia cross-talk and immune microenvironment dynamics in cancer. Although our *in silico* spatial perturbation experiments are in the proof-of-concept stage, they show promising potential as a framework for generating testable hypotheses and identifying candidate molecular drivers of spatial organization and cell-cell dependencies.

We recognize several limitations of this work. While Steamboat’s center cell and environment metagenes are interpretable as reflecting prerequisites and consequences of intercellular relationships, respectively, they reveal associations rather than causality. Additionally, other intercellular dependencies – such as those driven by metabolite production or nutrient competition – may influence the observed patterns and warrant further modeling. Steamboat also requires single-cell resolution, motivating future extension to accommodate spot-based spatial transcriptomics.

Steamboat’s novel design has enabled unique analyses, opening new avenues for understanding spatial cellular patterns in tissues for a wide range of biological contexts in health and disease. Intermediate spatial scales – between local and global – are also supported in Steamboat’s implementation and may be worth further exploration, but need additional experiments and validation. Future efforts in data integration across diverse modalities and technologies will also be crucial to maximize Steam-boat’s applicability. Unlike recent attention-based models that focus on linking genes [46, 47, 59, 60], Steamboat emphasizes relationships between cells. Looking ahead, we anticipate that this model will support more efficient and scalable representations of spatial cell states – an essential step toward building cellular foundation models capable of capturing organization and function at multiple scales [61].

## Methods

### The Steamboat model

#### Input data

Steamboat takes as input one or more spatial transcriptome (or other molecular profile) slides (see **Fig. 1**). When multiples slides are provided, the number of cells per slide may vary, but the gene set is assumed to be consistent across all slides. For a given slide with *N* cells and *G* genes, cell coordinates are preprocessed into a spatial neighbors graph *S* = (*V, E*) using *k*-NN method (default *k* = 8). The normalized and log-transformed expression profile is represented as a nonnegative real-valued matrix 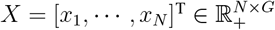, which is then fed into the attention model in Steamboat.

#### Introduction to the attention mechansim

The attention mechanism [32], originally introduced in natural language processing, enables each element (e.g., a word) to be understood in the context of surrounding elements. In Steamboat, this idea is further developed for spatial omics, where each “word” becomes a cell, and attention models contextual relationships between cells, enabling subsequent context-aware modeling. This is realized by a module called “attention head”. Each attention head is designed to recognize a specific type of relationship. For example, in language, one head might link nouns to adjectives, while another links verbs to their subjects. Similarly, in tissue, heads may attend to spatially or functionally related cells.

Mathematically, each element (i.e., word) is initially represented by a context-agnostic vector *x*. Each head uses two transformation matrices to link related elements: query *W* ^(q)^ and key *W* ^(k)^. For each element, the head calculates its “query vector” 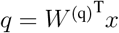. The “key vector” is also calculated for every element in the context 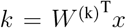. By taking the inner product of these vectors *q*^T^*k* (and after necessary normalization), the attention head assigns every pair of elements an attention score *a*, which reflects how closely related two elements are to each other. In other words, it matches the “query” with the appropriate “keys”.

Because cells also interact in different ways and strengths, Steamboat modifies this framework in several ways for biological interpretability and efficiency. Above all, our novel multiscale attention is tailored for cellular interactions of different distances. Moreover, all weights are constrained to be non-negative. Finally, query and key vectors are scalars, and their transformation matrices are single-column vectors. These vectors are called metagenes, as they represent weighted combinations of genes.

#### *Hyperparameters and trainable parameters in* Steamboat

Steamboat consists of *H* attention *heads* (a hyperparameter), each containing a set of metagenes, mathematically represented as vectors in gene expression space. For each attention head *h*, three types of metagenes model a center cell and its surrounding environment:

- *center cell metagene* 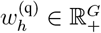 selects center cells;
- *environment metagenes* represent signatures of environments at three spatial scales: ego 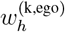, local 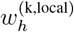, and global 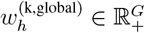;
- *reconstruction metagene* 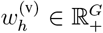 reconstructs the center cell based on its own features and the environment.

Since the ego environment refers to the center cell itself, its environment metagene 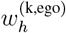 shares the same direction as the center cell metagene, but is scaled by a nonnegative scalar *κ*_*h*_ ∈ R_+_ to adjust its magnitude. Conceptually, the environment sends a signal (key), the center cell receives it (query) and responds with a reconstruction metagene. However, it is important to note that Steamboat operates based on correlation, and the causal directionality of these interactions remains an area for future development. A summary of all trainable parameters is provided in **Table** 1.

**Table 1.** Trainable metagene weights in Steamboat.

| Name | Notation |
| --- | --- |
| Center cell metagene | $w_h^{(q)} \in \mathbb{R}_+^G$ |
| Ego environment metagene (depend on $w_h^{(q)}$ and $\kappa_h$) | $w_h^{(k,ego)} := \kappa_h w_h^{(q)} \in \mathbb{R}_+^G$ |
| Local environment metagene | $w_h^{(k,local)} \in \mathbb{R}_+^G$ |
| Global environment metagene | $w_h^{(k,global)} \in \mathbb{R}_+^G$ |
| Reconstruction metagene | $w_h^{(v)} \in \mathbb{R}_+^G$ |
| Ego attention weight (scalar) | $\kappa_h \in \mathbb{R}_+$ |

#### *Attention mechanism in* Steamboat

The interaction between metagenes in Steamboat follows a modified multi-head attention mechanism [32] (**Fig. 1b**). Consider a center cell 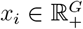 and its neighborhood cells *x*_*j*_ ∈ ***𝒩*** (*i*). Each center cell is assigned a *center cell score* indicating its association with a specific attention head *h*:

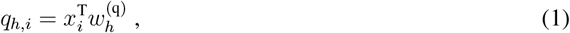

Cells in the ego, local, and global environments receive scores from their respective metagenes. The ego *environmental score* is:

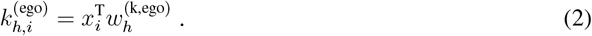

The *raw ego attention score*, which captures the interaction strength within the ego environment, is given by:

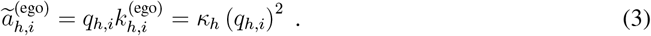

For local attention, each neighboring cell *x*_*j*_ is assigned an *environment score*:

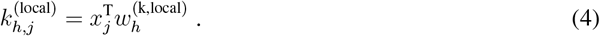

The *pairwise raw local attention score* between a center cell *x*_*i*_ and a neighbor *x*_*j*_ is:

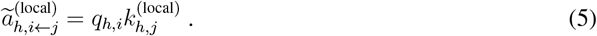

The *total raw local attention score* for cell *x*_*i*_ is computed as the average over its pairwise scores:

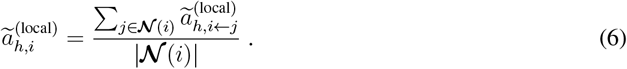

For global attention, the environment expands to the entire sample (*j* = 1 … *N*). The *global environment score* is:

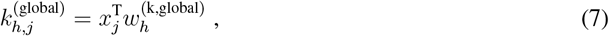

which defines the *pairwise raw global attention score*:

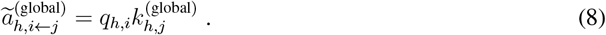

The *total raw global attention score* is then:

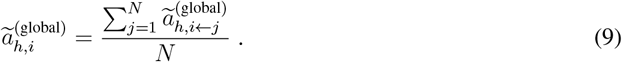

The three scales (ego, local, and global) follow the same attention mechanism structure but differ in the neighborhood definitions. The ego scale uses the center cell metagene as the environment metagene, as the sender and the receiver are the same cell. The local neighborhood excludes the center cell, while the global neighborhood includes both the center cell and the local neighborhood. This design maintains biological relevance while improving computational efficiency.

To avoid confusion, we use “score” for the result of *q*(·), *k*^(*…*)^(·), and *a*(·) for a cell, while “loading” refers to the gene weights in the metagenes *w*^(q)^, *w*^(k,*…*)^, and *w*^(v)^.

Once the total raw attention scores are computed for all heads, they are normalized to produce the final *attention scores* for each cell (scale ∈ {ego, local, global}):

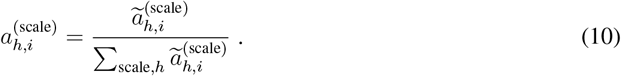

The *embedding* of a cell is an *H*-dimensional vector constructed by summing attention scores across scales:.

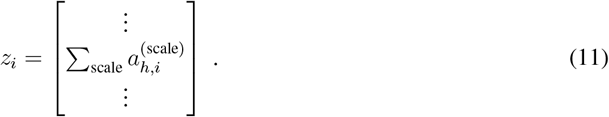

This embedding is then used to reconstruct the expression profile (*the reconstructed cell*) using the *value metagenes w*^(v)^:

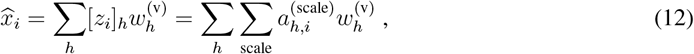

where [·]_*h*_ denotes the *h*-th entry of a vector. Because both *X* and the metagenes are nonnegative, all raw attention scores remain nonnegative, allowing simple normalization instead of a softmax function.

This structure resembles the multi-head attention mechanism in transformer [32], using masked attention with a single hidden dimension (i.e., *k* and *q*) to determine attention weights, along with a projection vector (*v*) for output transformation. To enhance interpretability, all weight vectors are constrained to be nonnegative. This eliminates ambiguity about why a cell attends to another, which can occur in previous models where positive and negative weights make attention head behavior difficult to interpret. Steam-boat’s constraints ensure that attention mechanisms remain interpretable and biologically meaningful.

#### Training strategy

The Steamboat model is trained using a self-supervised mask-and-predict approach [62], optimized with mean-squared error (MSE) loss and the Adam optimizer [63]. If multiple samples are available, batch gradient descent is applied by summarizing gradients obtained from each sample. The implementation includes sparse graph handling to improve both time and memory efficiency.

#### Number of attention heads

The number of attention heads, *H*, is a hyperparameter. We use PCA as a surrogate and choose *H* such that *H* principal components yield sufficiently clear cell clusters in UMAP.

#### Computational complexity

On 2.6 million cells across 129 slides (processed in mini-batches), Steamboat converges in 800 epochs, taking under one hour on a desktop computer with a Ryzen 5 3600 CPU (6 cores, 3.6 GHz) and an RTX 3080 GPU. The downstream Leiden clustering takes approximately three hours, similar to BANKSY [30], SEDR [20], and STAGATE [28].

### Interpreting Steamboat metagenes and embeddings

The key outputs of Steamboat are the cell embedding, reconstructed cell, and the weighted attention graph of cells (details in “Spatial domain segmentation” below) (**Fig. 1b**), which can be used for various downstream analyses (**Fig. 1c**).

#### Gene set enrichment

To measure the level of agreement of Steamboat metagenes and known gene sets, we use the AUROC (area under the receiver operating characteristic curve), inspired by the AUCell score [64]. We formulate a binary classification task: use the scores for genes in a metagene to predict whether a gene belongs to a given gene set. Specifically, we choose all possible cutoffs for the gene scores and calculate the TPR (true positive rate; range [0, 1]) and FPR (false positive rate; range [0, 1]), forming the ROC curve. The area under the curve is the AUROC. The range of AUROC is [0, 1], where values*>* 0.5 indicate positive enrichment and *<* 0.5 indicate negative enrichment.

#### Cell type clustering and visualization

The cell embeddings are used for cell type clustering and visualization. For clustering, we generate a *k*-NN *similarity graph* from these embeddings and apply Leiden clustering. For visualization, UMAP [65] is applied to the *k*-NN graph.

#### Spatial domain segmentation

Spatial domains are segmentations of a spatial omics slide, typically comprising regions that contain several distinct cell types or states [30], often with biologically meaningful interactions [29]. To capture these interactions, we generate a weighted attention graph for each local head, where edge weights are defined by the pairwise raw local attention score (**Eqn. 5**), normalized across all attention heads (using the same normalization factor as in **Eqn. 10**):

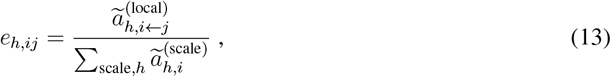

which forms the attention maps/matrices *A*_*h*_ = [*e*_*h,ij*_]. Since each attention graph is a weighted version of the spatial *k*-NN graph, all graphs share the same underlying topology (i.e., the sparse matrix representations share nonnegative entries). To aggregate them into one graph, we sum the edge weights across all heads (*A* = ∑_*h*_ *A*_*h*_). This graph is propagated *η* times using the transformation 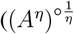, where “∘” denotes elementwise operation; default: *η* = 3) to reduce fragmentation of spatial domains.

To further integrate distinct cell type/state information, we construct another *k*-NN graph based on the degrees (i.e., the sum of edge weights) of nodes in the weighted attention graphs ([∑_*j*_ *e*_*h,ij*_]). Finally, we combine the propagated spatial *k*-NN graph and the cell type/state *k*-NN graph by summation, and apply the Leiden algorithm to segment the resulting graph into spatial domains.

#### Weight of heads and environment scales

Each attention head in Steamboat independently allocates attention across ego, local, and global scales. The weight of each scale is proportional to the norm of its corresponding environment metagenes, helping to identify which cells (and genes) are most influenced by their environment and through what mechanisms. The importance of each head is assessed post hoc by summarizing its attention score across cells.

#### Spatial perturbation

Spatial perturbation is a key capability enabled by Steamboat. We explore two types of perturbation. (1) Transplantation: We investigate how cells respond to being placed in a different spatial environment. This is implemented by moving cells of a given type to a new spatial region. Their expression profiles are then predicted using the model, and DEGs are identified by comparing relocated cells to those in the original location. (2) Gene knockout: We simulate a knockout of a gene in a specific cell type and assess how neighboring cells would respond. This is done by setting the expression of a particular gene in the target cell type to zero, then reconstructing the expression profile of surrounding cells. The most affected neighboring cells and their DEGs are identified.

#### Ligand-receptor interaction recognition

Ligand-receptor interactions are a fundamental mechanism of spatial cellular communication. To determine whether a local attention head corresponds to known ligand-receptor signaling, we reference curated ligand-receptor databases such as CellChatDB [51] and apply a permutation test to detect en-riched ligand-receptor pairs in the key and query metagenes. Specifically, we randomly shuffle metagene loadings to generate a null distribution, and assign empirical *P*-values to known ligand-receptor pairs.

#### Sample-level characterization

Global attention heads generate per-sample global key scores that summarize sample-level, macroenvironmental features. These scores can be correlated with phenotype or clinical variables, such as patient survival. The associated metagenes may reveal relevant signaling pathways. Survival analysis is performed using the Kaplan-Meier estimator, and differences between groups (e.g., CLR vs. DII; high vs. low global attention) are calculated using the log-rank test.

Overall, Steamboat leverages underlying cell-cell interactions – rather than simply smoothing across space – to construct cell embeddings and cellular interaction graphs. This approach enables unified and interpretable cell type clustering, spatial domain segmentation, and additional analyses such as ligand-receptor inference, sample classification, and spatial perturbation.

### Related methods and benchmark metrics

We evaluated Steamboat by comparing it with the following methods for cell-type clustering and/or spatial domain segmentation. (1) BANKSY augments each cell’s expression profile with neighborhood information [30], using a single hyperparameter *λ* to control spatial smoothing strength. In contrast, Steamboat treats clustering and segmentation as related but distinct tasks, solving via two outputs from a single model. Following the original publication of BANKSY, we set *λ* = 0.2 for cell type clustering and *λ* = 0.8 for spatial domain segmentation. Clustering resolution was tuned via grid search to match the ground truth number of clusters, with PCA dimensions set to 30 (which outperformed the default of 20). (2) STAGATE applies a graph attention autoencoder to incorporate neighborhood information prior to domain segmentation [28]. Like other graph-based networks, it performs inherent smoothing [31]. We used STAGATE via the DANCE framework [66], with the true number of spatial domains provided (a favorable setup for STAGATE). All other parameters were default. (3) SEDR applies a deep autoencoder gene expression embedding, combined with a variational graph autoencoder to capture spatial structure [29]. Because graph reconstruction is part of the training objective, the resulting embeddings are spatially smoothed. The ground-truth number of spatial domains was also provided. The number of neighbors was set to eight, which performed better than the default of six.

Steamboat was run with default parameters: *k* = 8 for the *k*-NN spatial graph and *k* = 15 (default in Scanpy) for the similarity graph. The training masking rate was set to 0.1. Leiden clustering resolution was set to 0.8 for spatial domain segmentation and 1.0 for clustering, tuned to approximate the ground-truth number of clusters.

For benchmarking, we used two commonly adopted metrics: (1) Adjusted Rand index (ARI) – better suited for clusters of similar sizes; (2) normalized mutual information (NMI) – more robust to unbalanced clusters better [67]. In practice, both metrics showed strong agreement across evaluations and were applied to both cell type clustering and spatial domain segmentation.

## Code Availability

The source code of Steamboat can be accessed at: https://github.com/ma-compbio/Steamboat.

## Acknowledgements

This work was supported, in part, by National Institutes of Health Common Fund 4D Nucleome Program grant UM1HG011593 (J.M.); National Institutes of Health Common Fund Cellular Senescence Network Program grant UG3CA268202 (J.M.); and National Institutes of Health grants R01HG007352 (J.M.), R01HG012303 (J.M.), R21DA061481 (J.M.), and U24HG012070 (J.M.). J.M. was additionally supported by the Ray and Stephanie Lane Professorship, a Guggenheim Fellowship from the John Simon Guggenheim Memorial Foundation, a Google Research Award, and a Single-Cell Biology Data Insights award from the Chan Zuckerberg Initiative. S.L. is a Lane Fellow. The funders had no role in study design, data collection and analysis, decision to publish or preparation of the manuscript.

## Author Contributions

Conceptualization: S.L. and J.M.; Software: S.L.; Investigation: S.L., J.T., G.W., and J.M.; Writing: S.L., J.T. and J.M. Funding Acquisition: J.M.

