## Supplemental Information for "Steamboat: Attention-based multiscale delineation of cellular interactions in tissues"

#### Supplementary Figures

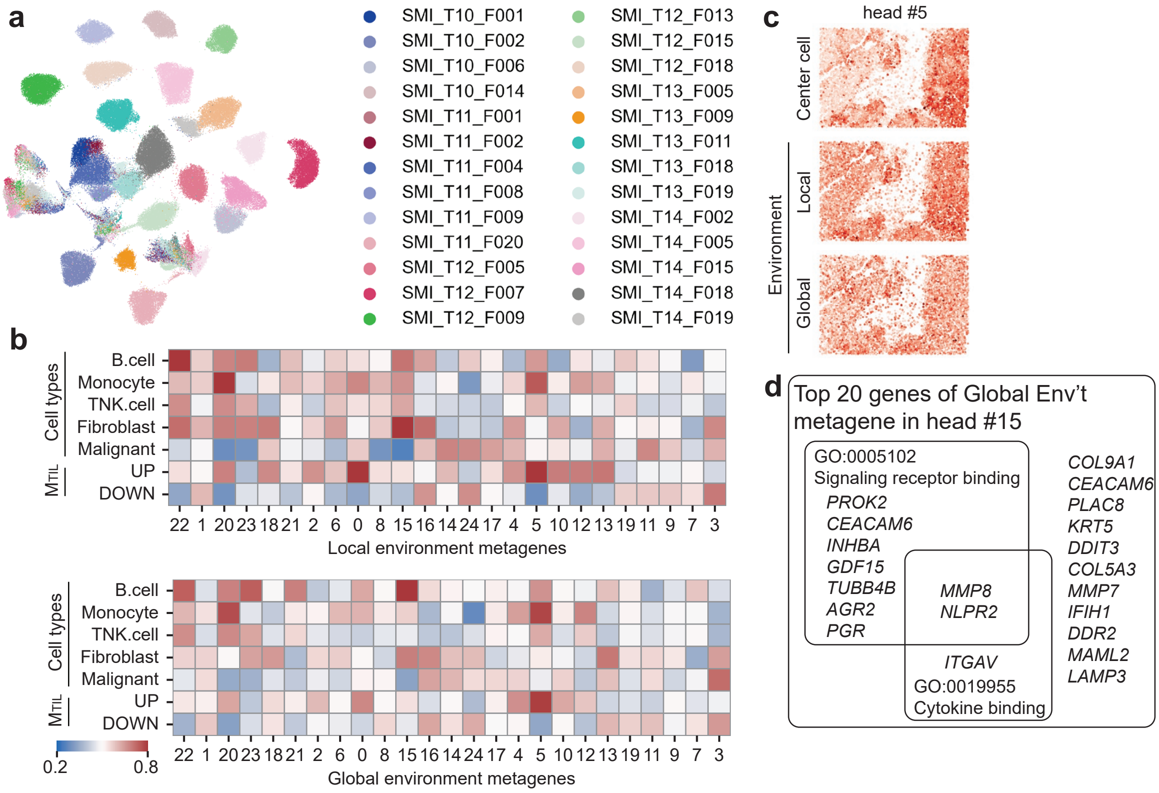


**Figure S1: Additional results from STEAMBOAT analysis of the ovarian cancer dataset. a.** UMAP visualization of STEAMBOAT embeddings, colored by samples. **b.** Gene set enrichment of local and global environment metagenes. **c.** Spatial distribution of cells selected by STEAMBOAT heads. **d.** Top 20 genes in the global environment metagene of head #15, grouped by gene ontology category.

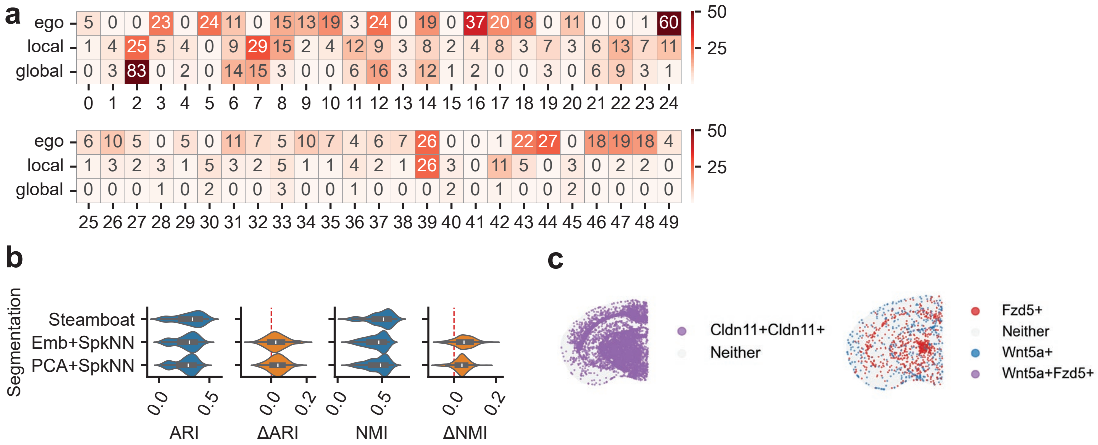


**Figure S2: Additional results from STEAMBOAT analysis of the mouse brain dataset. a.** Importance of the three attention scales across attention heads. Top: normalized by each head. Bottom: normalized over all heads. **b.** Ablation study results for spatial domain segmentation. **c.** Spatial expression distribution of *Cldn11*, *Fzd5*, and *Wnt5a*.

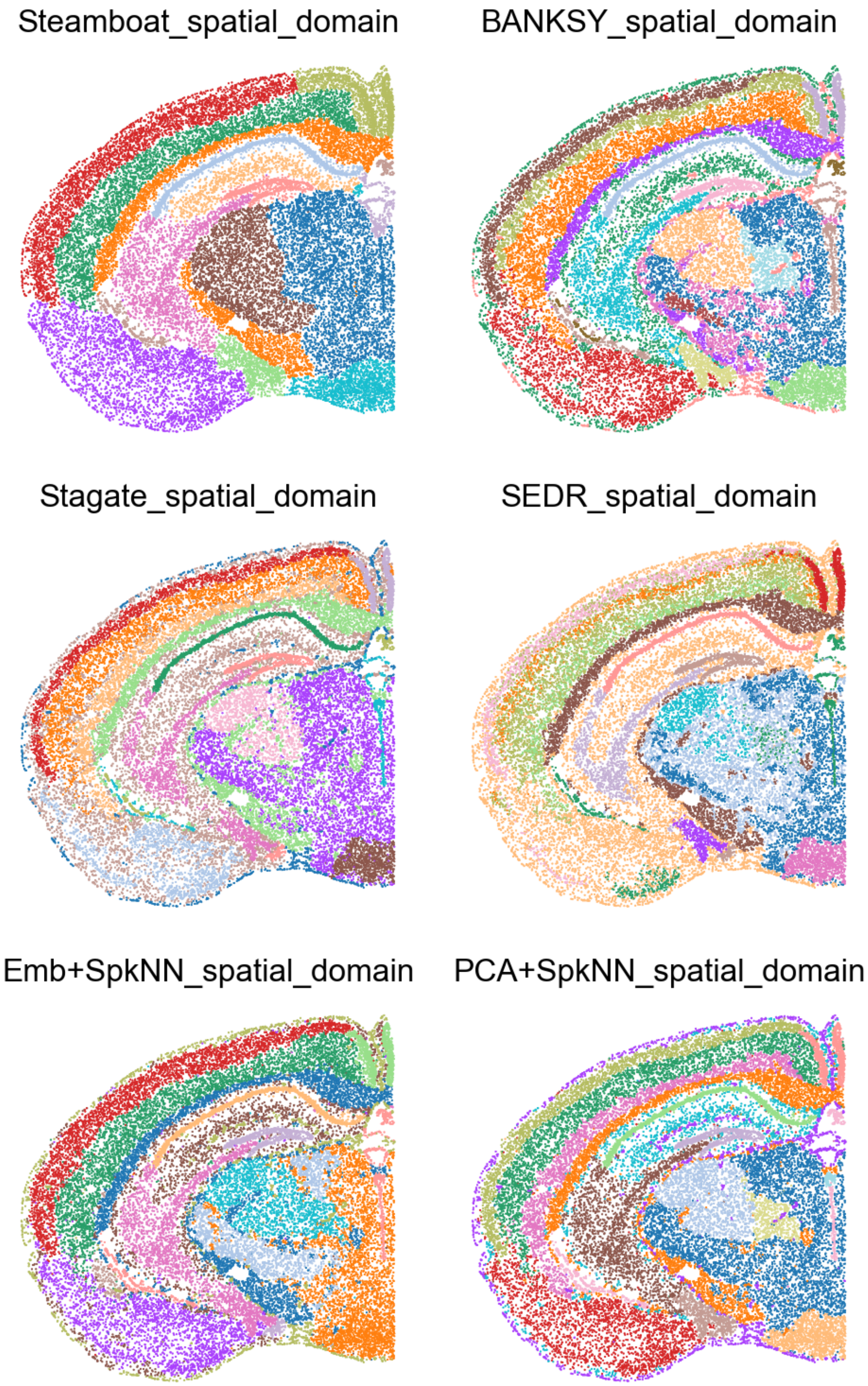


**Figure S3:** Spatial domain segmentation results from compared methods in the mouse brain dataset. See **Fig. 4**b for ground truth spatial domain annotations.

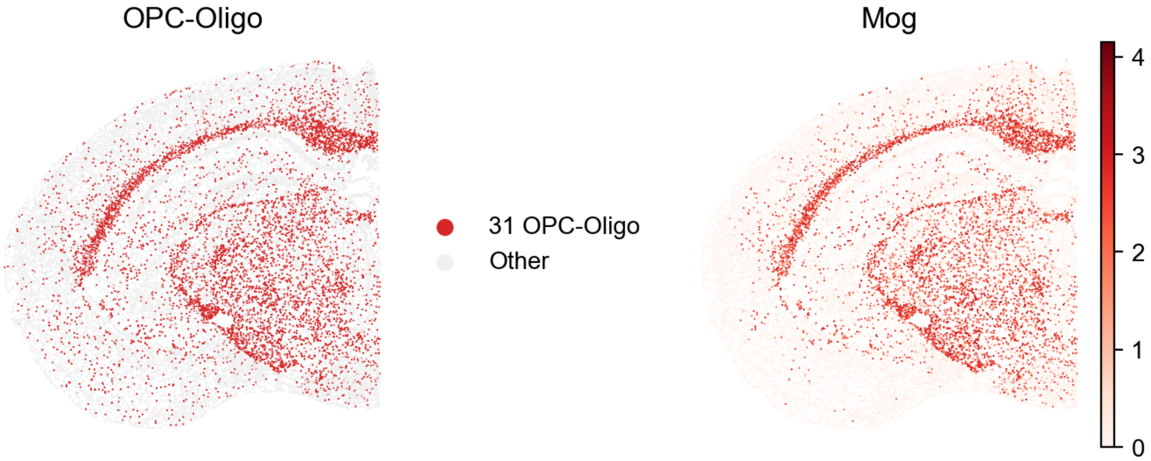


**Figure S4:** Spatial distribution of oligodendrocytes and OPC-Oligo cells is highly correlated with the expression of *Mog*.

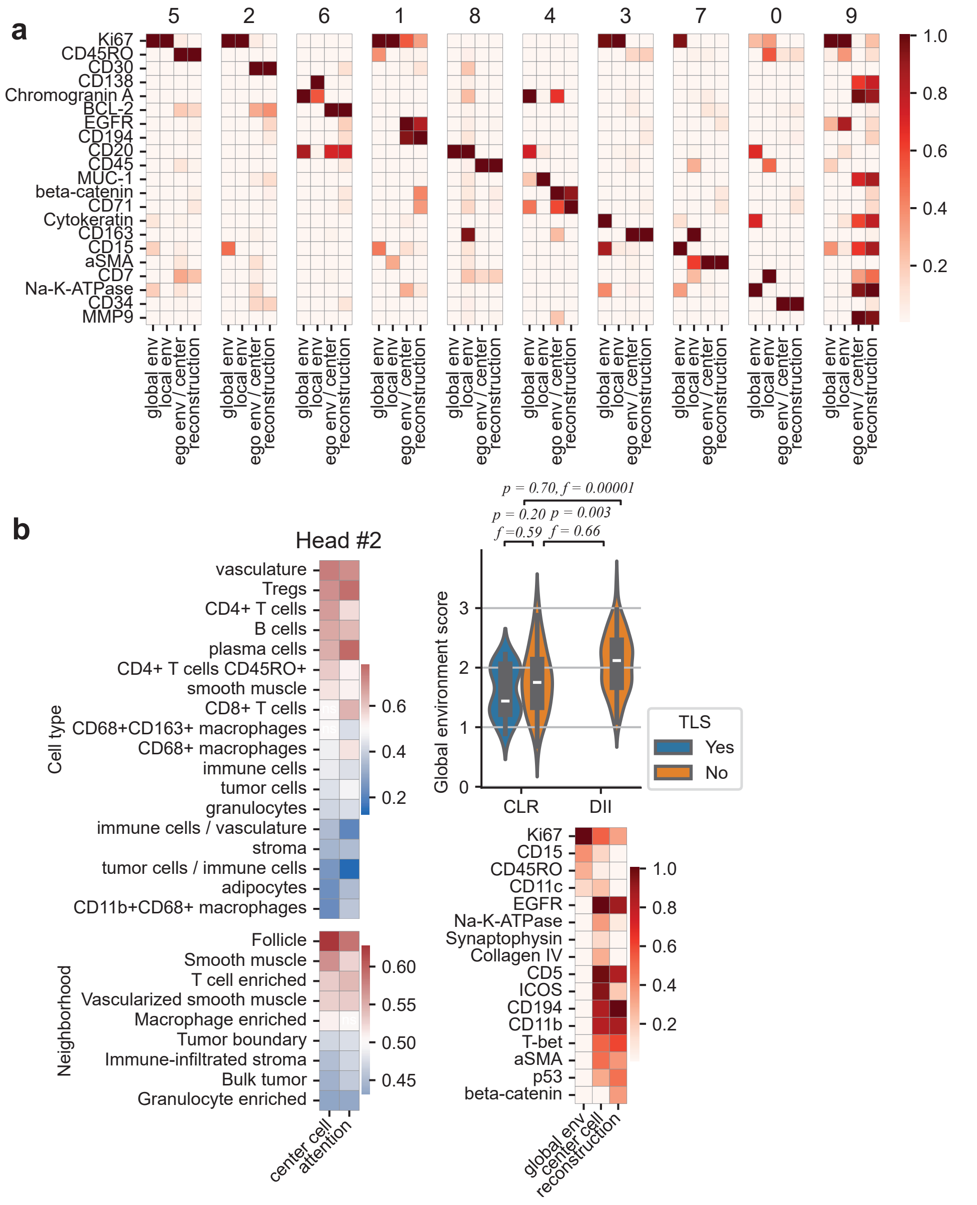


**Figure S5: Additional results from STEAMBOAT analysis of the colorectal cancer dataset. a.** Top genes in the metagenes of all attention heads. **b.** Characterization of head #2, including cell type and niche enrichment, metagene loadings, and association with CLR/DII classification.

### Supplementary Notes

#### Abbreviations of cell types in the mouse brain dataset

Abbreviations in **Fig.** 4 are listed as follows.

- Astro: astrocyte
- CB: cerebellum
- CGE: caudal ganglionic eminence
- CNU: cerebral nuclei
- CTX: cerebral cortex
- CTXsp: cortical subplate
- DG: dentate gyrus
- Epen: ependymal
- EPI: epithalamus
- ET: extratelencephalic
- GC: granule cell
- HB: hindbrain
- HPF: hippocampal formation
- HY: hypothalamus
- IMN: immature neurons
- IT: intratelencephalic
- LGE: lateral ganglionic eminence
- MB: midbrain
- MGE: medial ganglionic eminence
- MM: medial mammillary nucleus
- OB: olfactory bulb
- OLF: olfactory areas
- Oligo: oligodendrocytes
- OPC: oligodendrocyte precursor cells
- P: pons
- TH: thalamus
- V3: third ventricle
- VL: lateral ventricle
- Dopa: dopaminergic
- GABA: GABAergic
- Glut: glutamatergic
